# Rates of primary production in groundwater rival those in oligotrophic marine systems

**DOI:** 10.1101/2021.10.13.464073

**Authors:** Will A. Overholt, Susan Trumbore, Xiaomei Xu, Till L.V. Bornemann, Alexander J. Probst, Markus Krüger, Martina Herrmann, Bo Thamdrup, Laura Bristow, Martin Taubert, Valérie F. Schwab, Martin Hölzer, Manja Marz, Kirsten Küsel

## Abstract

The terrestrial subsurface contains nearly all of Earth’s freshwater reserves^1^ and harbors upwards of 60% of our planet’s total prokaryotic biomass^2,3^. While genetic surveys suggest these organisms rely on *in situ* carbon fixation, rather than the translocation of photosynthetically derived organic carbon^4–6^, corroborating measurements of carbon fixation in the subsurface are absent. Using a novel ultra-low level ^14^C-labeling technique, we show that *in situ* carbon fixation rates in a carbonate aquifer reached 10% of the median rates measured in oligotrophic marine surface waters, and were up to six-fold greater than those observed in lower euphotic zone waters where deep chlorophyll levels peak. Empirical carbon fixation rates were substantiated by both nitrification and anammox rate data. Metagenomic analyses revealed a remarkable abundance of putative chemolithoautotrophic members of an uncharacterized order of Nitrospiria – the first representatives of this class expected to fix carbon via the Wood-Ljungdahl pathway. Based on these fixation rates, we extrapolate global primary production in carbonate groundwaters to be 0.11 Pg of carbon per year.

## Main

The continental subsurface is the planet’s largest carbon reservoir^7^, housing up to 19% of its total biomass^2,8^ and 99% of its freshwater^1^. Despite accounting for only 6% of total stores, modern groundwater, *i*.*e*., the fraction accrued in aquifers over the past 50 years, is the single-most significant source of potable water. Carbonate karst aquifers alone are thought to supply people with nearly 10% of their drinking water^9^. Unfortunately, modern groundwater is also the most vulnerable to anthropogenic and climatic impacts^1^. While subsurface ecosystems have long fascinated ecologists^10^, and more recently microbiologists^11^, accessibility, enormous spatial heterogeneity, and the complete lack of process rate measurements has obscured a meaningful understanding of their contributions to global biogeochemical cycles^12^.

The widespread recognition that Earth’s biosphere extends deep into the subsurface occurred only recently^13^. Historically, carbon supply in such environments was thought to be limited to the trickling of surface-produced organic matter into the shallow subsurface^5^, or what was stored within sedimentary rocks^14^. In stark contrast, recent studies have shown that millimolar concentrations of dissolved H_2_ amassed in deep Precambrian Shield fracture groundwaters support the proliferation of chemolithoautotrophs^15^ and ultimately bacterivore nematodes^16^. A wealth of compelling genetic evidence suggests that *in situ* carbon fixation is critical for sustaining highly-diverse microbial metabolic networks in groundwater, both in the shallow and deep subsurface^4,17–23^. Despite the implications of gene-based surveys, the empirically derived activity measurements required to corroborate such inferences, constrain biogeochemical fluxes, understand system dynamics, and integrate processes into regional and global models have yet to be reported. Here we report our use of a novel radiocarbon method to derive empirical carbon fixation rates and place them in the context of global groundwater.

### Groundwater carbon fixation rates resemble those of marine surface waters

In this first ever evaluation of primary productivity in the shallow subsurface (*i*.*e*., groundwater wells 5 – 90 m deep), experimental carbon fixation rates varied from 0.043 ± 0.01 to 0.23 ± 0.10 μg C l^-1^ d^-1^ (mean ± SD; Fig. 1 A, Table S2, Supp. Info.). The ultra-low level ^14^C labeling approach developed in this investigation exploits the high sensitivity of accelerator mass spectrometry, thereby minimizing impacts to groundwater hydrochemical equilibria and affording shorter incubation times. This method is particularly useful within a carbonate geological setting, where high dissolved inorganic carbon (DIC) backgrounds and a scarcity of microbes warrant greater sensitivity than is achievable via scintillation-based ^14^C-labeling approaches. Rates resulting from our novel labeling technique likely approximate net primary productivity rather than gross productivity, as has been reported for marine systems^24,25^, and we further expect them to be conservative estimates for carbon fixation (Supp. Info.).

**Fig. 1.**
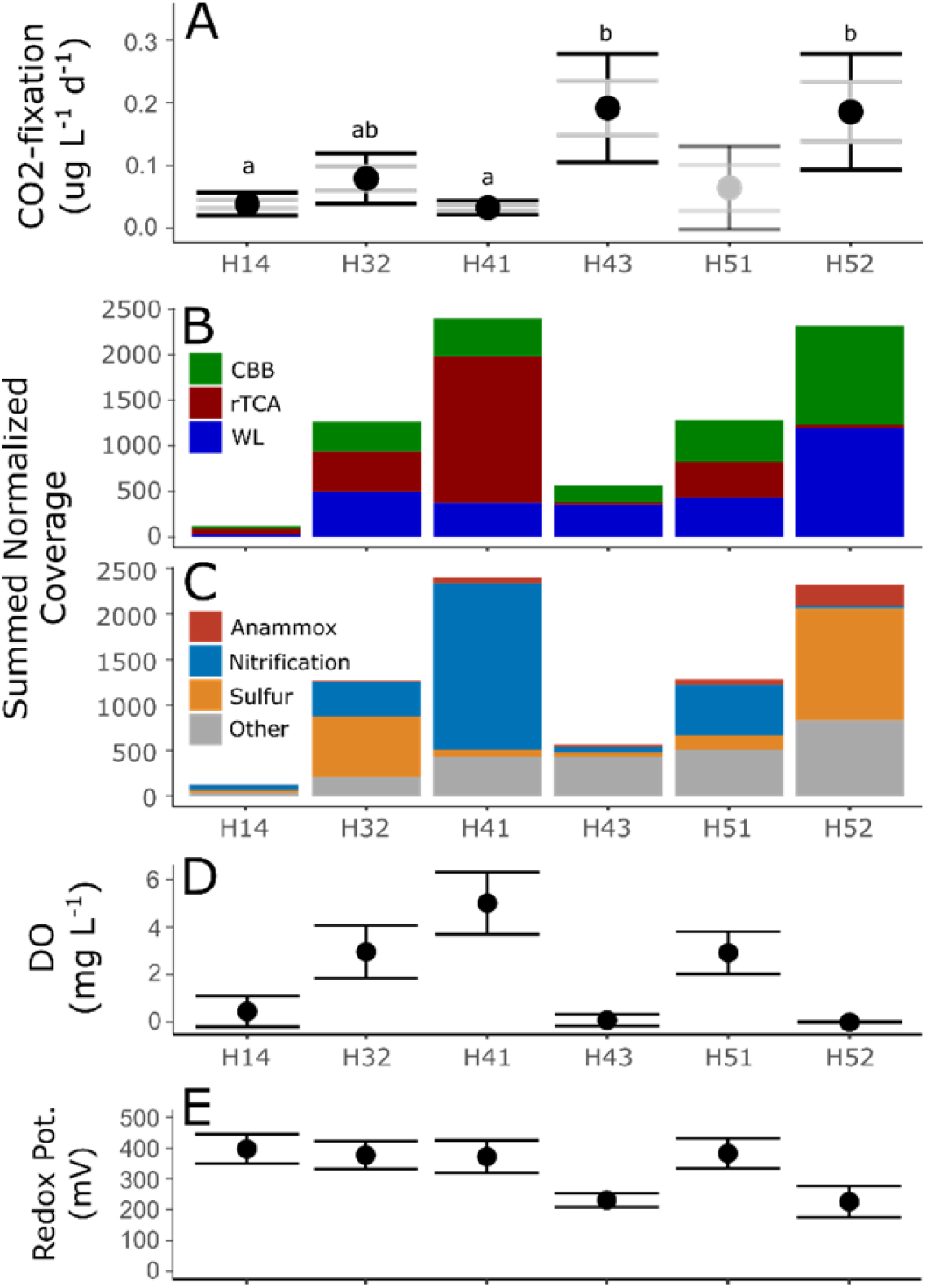
(A) Rates of carbon fixation across the aquifer transect. Outer error bars depict one standard deviation while inner grey bars delineate standard error of the mean. Rates for well H51 are derived from non-labeled controls (see Supp. Info.). Letters denote the results of ANOVA and posthoc Tukey tests (H51 was excluded from testing). (B) Relative importance of the predicted carbon fixation pathways and (C) electron donor sources in each well. Values are averages from the triplicate 0.2-μm filtered fraction metagenome samples. (D) Mean dissolved oxygen concentrations in groundwater collected in summer months (May through September 2010 - 2018), error bars depict standard deviation. (E) Redox potential measurements from identical time points.

We compared these carbon fixation rates that were measured in groundwater of varying biogeochemical characteristics^26^ with the only other subsurface ^14^CO_2_ assimilation measurements reported: those of a deep (830-1078 m) groundwater borehole from crystalline bedrock in Sweden ^27^. To do so, we converted the published rates of isotopic incorporation to carbon equivalents, revealing the lower but overlapping range of 0.0095 to 0.056 μgC l^-1^ d^-1^.

To better understand the relevance of the rates measured, we further compared them to those of well documented oligotrophic marine surface waters. Unlike our samples, the carbon fixed in these waters was sourced almost entirely by bacterial photoautotrophs^28,29^. When compared directly to a comprehensive dataset compiled by Liang *et al*. ^30^, our rates overlapped with those of global marine waters at depths to 140 m, equating to roughly 10% of the reported global median for 0-20 m depths (2.65 μg l^-1^ d^-1^, interquartile range (IQR) = 1.74, 6.02), and 20% of the median for 20-140 m depths (1.2 μg l^-1^ d^-1^, IQR = 0.6, 1.7). Comparisons to the extensively studied Sargasso Sea in the Bermuda Atlantic Timeseries Study (BATS)^31,32^ and the Hawaiian Oceanographic Timeseries (HOTs)^33^ datasets yielded similar findings (Fig. 2). Our rate measurements ranged between three and 23% of the median reported net primary productivity in the upper euphotic zones (down to ∼ 120 m) and 20 to 600% of the median of the lower euphotic region (100 - 120m) where deep chlorophyll levels peak.

**Fig. 2.**
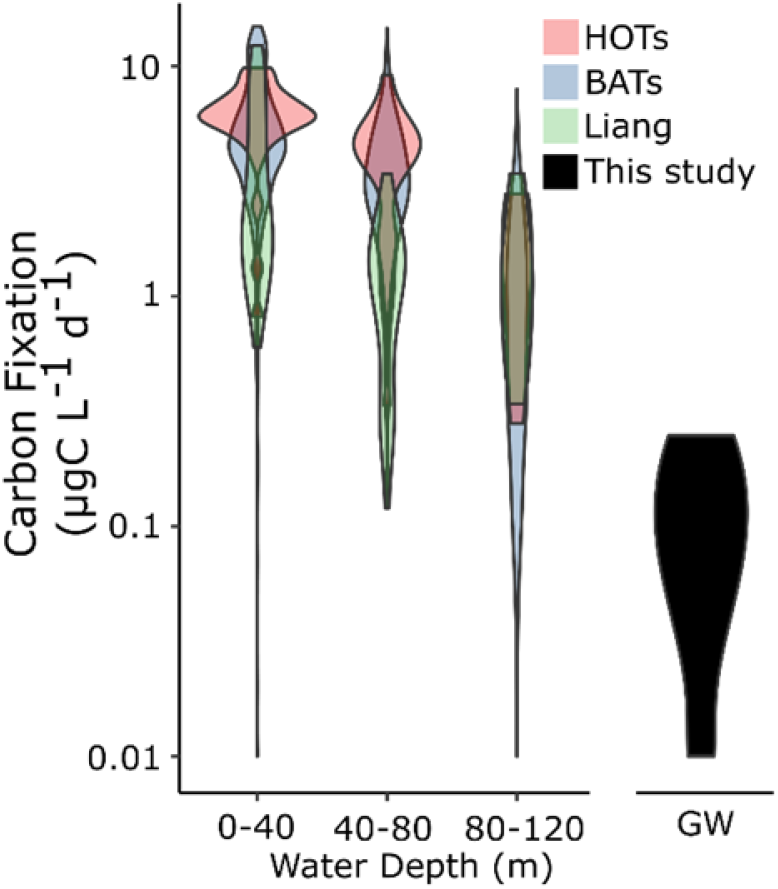
Violin plots depicting the distribution of carbon fixation rates measured in oligotrophic marine surface waters and groundwater. HOTS = Hawaiian Oceanographic Timeseries (1999, cruises 101-110), BATs = data from 1988-2016 for the Bermuda Atlantic Timeseries, and Liang = a collated dataset compiled by Liang *et al*., 2017. GW represents the range of groundwater samples within the Hainich CZE.

We also considered contributions to existing particulate organic carbon (POC) stocks and new carbon inputs per microbial cell count. After normalizing for estimated total bacterial cell numbers, groundwater yielded 0.3 - 10.8 fg of fixed carbon per bacterial cell per day (Table S5), which matched estimates of 0.25 - 12.1 fg C per bacterial cell per day across the marine photic zone (5 - 150 m). However, groundwater received new daily carbon inputs of only 0.47% ± 0.22% (mean ± SD, Table S5) of its existing POC, much lower than the marine system’s 2.6% ± 2.9% gain in the lower euphotic zone and 22% ± 18% at the surface ^34,35^. This disparity might stem from the larger recalcitrant fraction of particulate organic carbon in groundwater compared to oligotrophic oceans, which is supported by deviations in ^14^C and ^13^C signatures of total groundwater POC concentrations compared to lipid signatures of resident microbes ^36^.

### An ecosystem dominated by chemolithoautotrophs

To identify dominant microbial primary producers, a total of 4,175 metagenome-assembled genomes (MAGs) were generated from groundwater samples, 1,224 of which passed quality thresholds. Of these, 102 putative chemolithoautotrophs exhibiting at least 50% completion scores for carbon fixation pathways were identified (Fig. 3). Almost exclusively bacterial (101), these MAGs represented 17 distinct phyla, 21 classes, and 35 families (Fig. 3, Table S3). In some samples, up to 12% of metagenomic reads from a sample could be recruited to these chemolithoautotrophic MAGs (Supp. Info., Fig. S3). A single archaeal MAG of the family Nitrosopumilaceae encoded gene products for the 4-hydroxybutyrate 3-hydroxypropionate pathway and was not relatively abundant (< 5x max normalized coverage).

**Figure 3.**
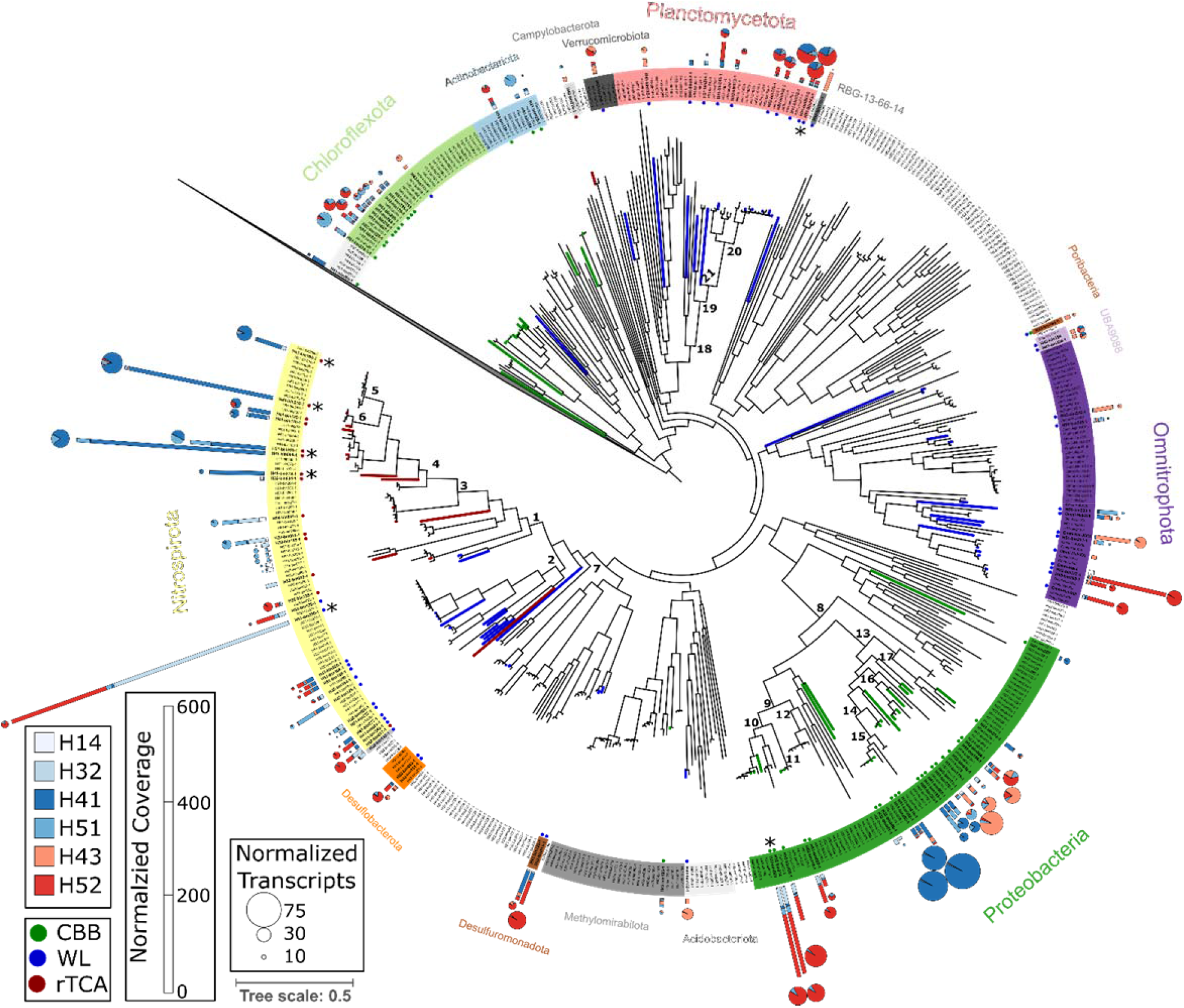
Approximately-maximum-likelihood phylogenetic tree based on concatenated single-copy protein alignments for all bacterial MAGs considered. Branches are colored according to the predicted carbon fixation pathway, and the matching leaf is indicated by a point. Bar charts present average normalized metagenomic coverages within each well from triplicate 0.2-μm filtered fractions. Pie charts show the coverage of mRNA transcripts recruited, normalized to gene number and library size. CBB = Calvin-Benson-Basham cycle, WL = Wood Ljungdahl pathway, rTCA = reverse TCA cycle. Asterisks denote MAGs discussed in greater detail in Supp. Info. The tree is rooted using Patescibacteria (CPR) as an outgroup, indicated by the collapsed grey leaf in the upper left. Node numbers represent: (1) c_Nitrospiria, (2) c_Thermodesulfovibrionia, (3) o_Nitrospirales, (4) f_Nitrospiraceae, (5) g_Nitrospira_D, (6) g_Nitrospira, (7) p_Nitrospinota, (8) c_Gammaproteobacteria, (9) o_Acidiferrobacterales, (10) f_Sulfurifustaceae, (11) g_SM1-46, (12) f_UBA6901, (13) o_Burkholderiales, (14 f_Nitrosomonadaceae, (15) g_Nitrosomonas, (16) f_SG8-41, (17) f_SG8-39, (18) c_Brocadiae (19) o_Brocadiales, (20) f_Brocadiaceae, and (21) f_Scalinduaceae.

Three chemolithoautotrophic pathways were detected (Fig. 1 B): the Calvin-Benson-Bassham (CBB), Wood-Ljungdahl (WL), and reverse TCA (rTCA) cycles were present in 37, 50, and 15 MAGs, respectively. The summed and normalized relative coverages of MAGs equipped with these metabolic pathways aligned with the carbon fixation rates measured in wells H52, H32, and H14, while contrasting with rate data from wells H41and H43 (Fig. 1, Supp. Info.). The greatest relative abundances of chemolithoautotrophs were detected in oxic well H41 and anoxic well H52. Anoxic groundwater was dominated by putative sulfur oxidizing (53% of summed and normalized coverages of all chemolithoautotrophic MAGs) and anaerobic ammonium oxidizing (anammox [10%]) autotrophic microbes, while oxic groundwaters harbored greater abundances of potential nitrifiers (76%; Fig. S4 B, Supp. Info.).

### Poorly characterized microbes influence carbon fixation potential

The most abundant putative chemolithoautotrophic populations represented by MAGs generated from anoxic groundwater were of poorly studied and/or uncharacterized microbial lineages. Those most abundant in oxic groundwaters, however, were phylogenetically and metabolically similar to well-characterized microbes (Supp. Info.). In both cases, metabolic reconstructions suggested that dominant subpopulations could access a diverse suite of (in)organic electron acceptors and donors. We mapped previously generated RNA-seq data ^37^ to these MAGs to confirm the active expression of gene products involved in energy acquisition and carbon fixation. As opposed to the broad distributions posited by DNA-based abundances, transcript data revealed far more restrictive ranges in which specific gene products were favored (Fig. 3). Given their metabolic versatility and the results of previous cultivation-based analyses ^38^, these populations are expected to be mixotrophic, *i*.*e*. capable of supplementing carbon requirements with available organic matter. Overall, carbon fixation in anoxic groundwater was predicted to be fueled by reduced sulfur and there were three highly abundant, sulfur-oxidizing MAGs identified, each accounting for > 2% of the total metagenomic reads in some samples (100-400x normalized coverages).

The most abundant MAG encountered in this study belongs to a deep-branching order, 9FT-COMBO-42-15, of class Nitrospira and is the first representative of Class Nitrospiria thought to fix carbon via the WL pathway (Fig. 3, Fig. 4A, Supp. Info.). As there is precedence for autotrophic WL-utilizing bacteria within phylum Nitrospirota, and *Ca*. Magnetobacterium was characterized with an equally flexible metabolism, these traits may be more widespread within the phylum than previously thought ^39^. In addition, two MAGs capable of coupling sulfur oxidation to carbon-fixation via the CBB cycle were identified as members of the Sulfurifustaceae family of Proteobacteria (Supplemental Information, Figure 4 B). These MAGs recruited 10-fold more transcripts than their Nitrospirota counterparts and were among the most transcriptionally active chemolithoautotrophic genomes detected (Fig. 3, Fig. 4, Supp. Info.). With its closest reference genomes *Sulfuricaulis limicola* and *Ca*. Muproteobacteria (RIFCSPHIGHO2_12_FULL_60_33), the taxonomic identity of this family is under debate. Per GTDB classification nomenclature, Muproteobacteria belong to the Sulfurifustaceae family, and members of this family have been posited to oxidize sulfur in both aquatic and terrestrial environments ^40–42^.

**Fig. 4.**
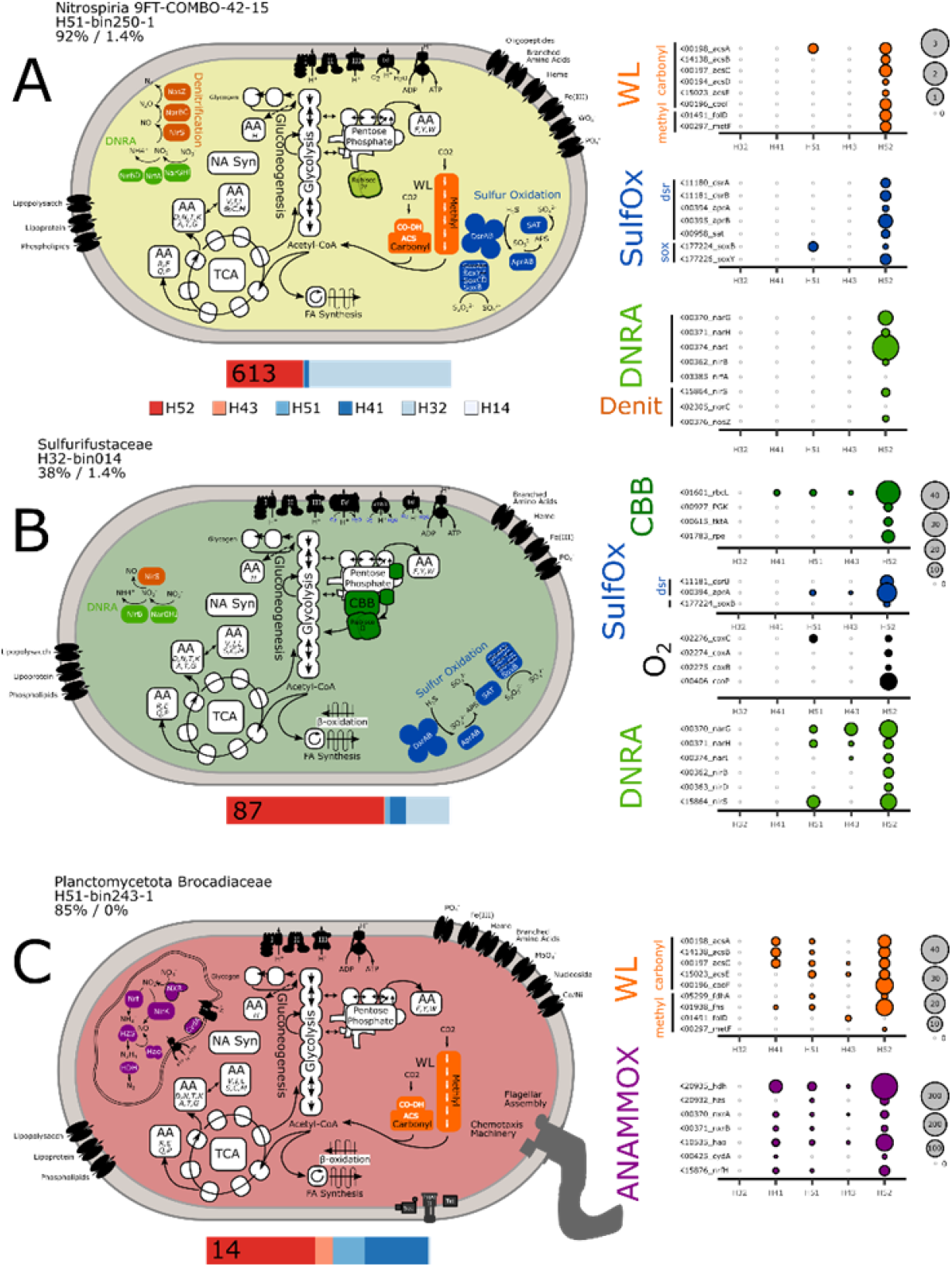
Metabolic reconstructions of dominant populations recovered from wells exhibiting both elevated rates of carbon fixation and relatively high abundances of MAGs representing chemolithoautotrophs. Bar charts below each metabolic model summarize the average normalized coverage across each sample, scaled proportionally. Values indicate the sum normalized coverage for each MAG. Balloon plots depict normalized transcript coverages for genes affiliated with each pathway. If multiple copies were present, only the most active copy was plotted.

Planctomycetota MAGs, predicted to couple anaerobic ammonium oxidation to carbon fixation via the WL pathway, exhibited mean transcriptional activities on par with their Sulfurifustaceae MAG counterparts (Fig.3, Fig. 4C, Supp. Info.). The elevated transcriptional activity of gene products germane to the CBB and WL pathways suggests that taxa wielding such functions play a disproportionately large role in chemolithoautotrophy relative to their DNA-based abundances. Surprisingly, all putative anammox MAGs detected were transcriptionally active in oxic groundwater (Fig. 3, Fig.4 B; wells H41 and H51). Anammox reactions are typically inhibited in the presence of oxygen ^43^, although microbes will still express critically important genes in low oxygen environments ^44,45^.

### Nitrogen-based rate measurements validate carbon fixation rates

To evaluate the relationship between anammox and carbon fixation in anoxic groundwaters, we compared the rates of each in a well harboring the greatest relative abundance of anaerobic ammonium oxidizing bacteria (well H52). Well H52 exhibited anammox rates of 1.2 ± 0.5 nmol N_2_ l^-1^ d^-1^. Empirical stoichiometric data demonstrates that 1.02 moles of N_2_ is produced via anaerobic ammonium oxidation for every 0.066 moles of CH_2_O_0.5_N_0.15_ reduced to biomass^46^. Assuming equivalent stoichiometry, the rate of carbon fixation via anammox in groundwater would be 0.93 ± 0.39 ng C l^-1^ d^-1^, more than 200 times lower than the 220 ng C l^-1^ d^-1^ measured. This result is corroborated by metagenomic data that suggest the high rate of carbon fixation in anoxic groundwater is more likely driven by reduced sulfur than reduced nitrogen.

Metagenomic and metatranscriptomic data predicted that nearly all of the organic carbon produced under oxic conditions in well H41 would be coupled to nitrification. To test this, we monitored the rate of aerobic ammonium oxidation in this well and recorded a mean production of 125.8 ± 5.9 nmol NO_2_^-^ + NO_3_^-^ l^-1^ d^-1^. Since the most abundant nitrifiers detected were most closely related to complete ammonium oxidizing bacteria (Supp. Info.), we based our calculations on the 394 mg protein per mol of ammonia growth yields of *Nitrospira inopinata*, a comammox organism^47^. Assuming a cellular composition of C_5_H_7_O_2_N^48^ and 55% protein content, we estimated a rate of 48.5 ± 1.9 ng C l^-1^ -d^-1^, which was well within the range of error for our measured rate of 43 ± 13 ng C l^-1^ d^-1^ confirming the importance of nitrification for carbon fixation at this site.

### Global estimates for groundwater primary productivity

There are an estimated 22.6 million km^3^ of groundwater on Earth ^1^, 2.26 and 12.66 million km^3^ of which are housed in carbonate and crystalline aquifers, respectively. If we assume that our average rates accurately represent carbonate groundwater systems, then 0.108 ± 0.069 Pg (mean ± SD) of carbon is fixed every year in this global ecosystem (Table S4). If the values reported from crystalline aquifers^27^ are representative of this environment, then another 0.15 ± 0.11 Pg C would be fixed there every year. Collectively, the net primary productivity of ∼ 66% of the planet’s groundwater reservoirs would total 0.26 Pg C yr^-1^, approximately 0.5% that of marine systems and 0.25% of global NPP estimates^49^. Applying the lowest measured values from each rock type yields the more conservative estimate of 0.079 Pg C yr^-1^, 0.076% of the global NPP. Although the total production rates in groundwater seem small, this is because of the relatively small volume of groundwater compared to the vast surface ocean - what is most surprising is that terrestrial subsurface fixation rates are approaching those of phototrophic organisms.

## Conclusions

Using a novel ultra-low level ^14^C-labeling technique to generate empirically derived estimates of primary productivity in groundwater for the first time, we showed that carbon fixation rates in a carbonate aquifer reached 10% of the median rates reported in oligotrophic marine surface waters and six-fold greater than those observed in the deep chlorophyll maximum of the lower euphotic zone. Normalizing rates according to estimated bacterial numbers revealed equivalent carbon input (*i*.*e*., 0.3 - 12 fg C per cell) for both systems, despite the fact that daily inputs of new POC were 40 times greater in marine waters than in groundwater. This disparity makes sensel, since trophic webs are simpler in the subsurface, and the export of organic matter is constrained by long water residence times within the aquifer. As the vast majority of photosynthetically derived carbon in marine systems is labile (half-life < 1 day), the findings of this study solicit new hypotheses regarding carbon cycling in the subsurface, particularly those positing newly synthesized carbon rather than surface-derived organic matter as the primary source of fuel for microbiota.

Complementary metagenomic analyses identified novel microbes capable of exploiting metabolic pathways previously unreported for their given phylotype. Carbon fixation rates coincided with the potential for chemolithoautotrophy in three of the five groundwater wells examined. Comprehensive metabolic reconstructions revealed versatile metabolisms with access to numerous sources of electron donors and acceptors, particularly in taxa detected in high abundance in anoxic and hypoxic groundwater. While these populations were widely distributed across broad biogeochemical regimes, the use of previously generated metatranscriptomes helped identify more specific activities.

Nitrogen-based transformations provided independent validation of carbon fixation rates in oxic waters and corroborated metagenomic data that hinted at the inconsequential impact of anammox on carbon fixation in anoxic groundwater. If our average rates accurately represent all carbonate groundwater systems, then 0.108 ± 0.069 (mean ± SD) Pg of carbon (0.22% of global marine NPP) is fixed every year under these geologic settings. Applying these rates of carbon fixation to ecosystem processes alters the way we think about these environments, challenges the importance of surface-derived organic matter fluxes on shallow subsurface functioning, and establishes a framework broadly applicable across groundwater systems.

## Methods

### Site description

Groundwater samples were sampled from the Hainich Critical Zone Exploratory (NW Thuringia, Germany) ^26,50,51^. This aquifer assemblage consists of a multistory, fractured system composed of alternating layers of limestone and mudstone that developed along a hillslope of Upper Muschelkalk bedrock ^26^. The primary aquifer, represented in this study by wells H41 and H51, is oxic and lies within the Trochitenkalk formation (moTK). Primarily suboxic to anoxic, mudstone-dominated overhanging strata lies within the Meissner formation (moM) and is represented here by wells H14 (moM - substory 1), H32 (moM - 5, 6, 7), H43 (moM - 8), and H52 (moM - 3, 4). Geochemically, H32 and H41 coalesce into a single cluster while each of the other wells represent distinct regimes. Consistent with previous microbiological characterizations, however, each well studied represented a distinct community state ^52^.

### ^14^C-DIC incorporation assay

This method, similar to a sensitive methane oxidation technique previously described^53^, is a modification of traditional ^14^C-CO_2_ primary productivity approaches^54^ predicated upon the sensitivity offered by accelerator-based mass spectrometry. Groundwater was collected in July 2020 during sampling campaign PNK130, as described by Herrmann et al.^23^. After approximately three well volumes had been discharged and physicochemical parameters stabilized, groundwater was collected directly into nine pre-sterilized 2-liter borosilicate bottles, from the bottom up. Bottles were then overfilled with > two volumes and sealed with gas-tight rubber stoppers. Triplicate samples from each well were then subjected to three treatments. A labelling treatment consisted of 6.77 × 10^−7^ mmol C-NaHCO_3_ which contained 200 Bq of activity [50 μCi; American Radiolabeled Chemicals (ARC), St. Louis, MO] diluted to 9.38 Bq/μl with sterilized milliQ water, adjusted to pH 10, and verified using a scintillation counter. An advantage of this ^14^C technique is that the small amount of tracer added (representing 0.000006% of the total DIC) did not change the substrate concentration or influence conditions like pH that could affect microbial populations. Kill controls were prepared in the same way, except 10 ml of 50% ZnCl_2_ (w/v; final conc. 36.7 mM) was added to inhibit microbial activity. Unamended groundwater was also used as a control. All bottles were incubated in the dark at near *in situ* temperature for ∼ 24 hours. Entire volumes were acidified to pH 4 with 3M HCl, bubbled with N_2_ for one h to remove DIC, and then filtered through pre-baked (550< 8 hours) quartz fibers (47 mm, 0.3um pore size, Macherey-Nagel QN-10) using pre-baked filter stands (EMD Millipore).

Filters were vacuum dried, sealed in quartz tubes with cupric oxide wire under vacuum, and combusted at 900 < for two hours. Evolved CO_2_ was purified cryogenically, measured as pressure in a known volume to determine C content, and reduced to graphite for measurement by accelerator mass spectrometry at the WM Keck Carbon Cycle Accelerator Mass Spectrometry facility^55^. From the label incorporation and amount of carbon retained on the filters (Supplemental Data File), fixation rates were calculated using equation (1):

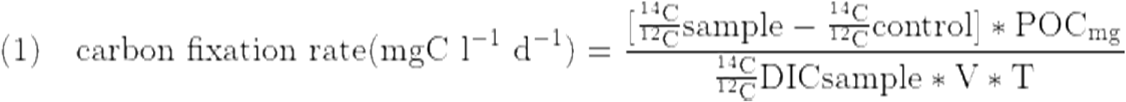

The technical variation was at most 3.6% (median = 0.78%) of the biological variation for the ^14^C measurements and was not considered in standard error of the mean calculations. Standard error of the mean was determined for both the ^14^C-based measurements (difference between two sets of triplicates, label and control, or label and kill controls) and POC measurements (all nine bottles from each well), separately. These errors were then propagated to yield the final error estimations. Analyses of variance and post-hoc Tukey HSD tests were conducted on resulting summary statistics (mean ± SEM) using the following utility: https://acetabulum.dk/cgi-bin/anova. All ^14^C enrichment values were calculated using the differences between the 200 Bq-labeled samples and the 200 Bq-labeled kill controls. Rates calculated based on no-label addition controls are presented in Table S2. Data from global oligotrophic marine systems was included from Supplementary Data Sheet 1^30^, the Bermuda Atlantic Timeseries years 1988-2016 via FTP (http://batsftp.bios.edu/BATS/production/bats_primary_production.txt) ^31^, and from the Hawaiian Oceanographic Timeseries via FTP (ftp://ftp.soest.hawaii.edu/hot/primary_production). POC data from both sites was extracted from Dryad datasets generated by Martiny et al. ^34,35^. Bacterial cell number estimates for HOTs were obtained from the FTP site: https://hahana.soest.hawaii.edu/FTP/hot/microscopy/EPIslides.txt.

### ^15^N-isotope incubation experiments

Groundwater from wells H41 and H52 was collected in September 2018 and November 2018 to measure nitrification rates and anammox rates, respectively. Briefly, groundwater was collected into sterile glass bottles, from the bottom up, using a sterile tube. Bottles were then overfilled with three volume exchanges, and sealed headspace free with silicone septa. Each sample was collected in triplicate alongside one control bottle per well. Samples were kept at 4°C until they were processed (no more than 2 h post-collection).

For nitrification measurements, 10 ml was removed from each sampling bottle (total vol. 0.5 L) and replaced with N_2_ to analyze inorganic nitrogen and pH. Groundwater from control bottles was sterile filtered through a 0.2 µm filter (Supor, Pall Corporation, USA). Sterile filtered ^15^N ammonium sulfate solution (98%, Cambridge Isotope Laboratories, Tewksbury), serving as a substrate for ammonia oxidizing prokaryotes, was then was added to a final conc. of 50 µM. Samples were incubated at 15°C in the dark *sans* agitation for five days. Ten-mL fractions were removed and replaced with N_2_ at the outset of the experiment and after 12, 24, 48, 70 and 120 hours via filtration through 0.2 µm filters, which were stored at −20°C for isotopic ratio mass spectrometry (IRMS) analyses. Additional 10 ml fractions were removed at intervals to monitor pH and inorganic nitrogen during the incubation.

For anammox rate measurements, sampling bottles (total vol. 1 L) were flushed with N_2_ under sterile conditions for 30 minutes to remove all remnants of oxygen. Five-mL fractions were removed and replaced with N_2_ from each sample (and control) bottle to assess background ^14^NH_4_^+^ concentrations. Subsequently, samples were spiked with either (a) 50 µM ^15^NH_4_ ^+^ + 5 µM ^14^NO_2_^-^ or (b) 5 µM ^15^NO_2_^-^ as previously described ^56^. Control bottles, serving as abiotic controls were sterile filtered (0.2 µm filters; Supor, Pall Corporation, USA) prior to flushing and the addition of nitrogen compounds. To facilitate destructive sampling at eight time points, groundwater (30 mL; in triplicate) was dispensed into sterile serum bottles leaving ∼ 8 mL of headspace. Bottles were immediately sealed with butyl septa, crimp sealed and the headspace was purged with He. All bottles were then incubated in the dark at 15°C *sans* agitation and incubations were terminated after 0, 12, 24, 36, 48, 60, 72 and 96 hours by adding 300 µl of 50% (v/w) aqueous zinc chloride solution.

Nitrification rates were determined based on ^15^NO_2_^-^ + ^15^NO_3_^-^ production in incubations with ^15^NH_4_ ^+. 15^NO_2_^-^ and ^15^NO_3_ ^-^ were converted to N_2_ via cadmium reduction followed by a sulfamic acid addition^57,58^. The N_2_ produced (^14^N^15^N and ^15^N^15^N) was analyzed on a gas chromatography isotope ratio mass spectrometer as previously described^59^. Rates were evaluated from the slope of the linear regression of ^15^N produced with time and corrected for the fraction of the NH_4_^+^ pool labelled in the initial substrate pool. The production of ^15^N labeled N_2_ from anammox was analyzed on the same IRMS as for nitrification rates, and calculated as described^60^. Note, denitrification was not detected in any of the ^15^NO_2_ ^-^ incubations. T-tests were applied (p < 0.05) to assess whether rates were significantly different from zero (Fig. S2).

### DNA extraction & sample preparation

Samples used to generate metagenomic libraries were collected in January 2019 during sampling campaign PNK 110. For each sample replicate, approximately 50 - 100 L of groundwater was filtered sequentially through 0.2 µm and 0.1 µm pore sized PTFE filters (142 mm, Omnipore Membrane, Merck Millipore, Germany; Table S1). With the exception of H32 (did not yield sufficient volumes), each well was sampled in triplicate. H32 was duplicated using a sample previously collected during campaign PNK108 (November 2018). Filters were frozen on dry ice and stored at −80 °C prior to extraction. DNA was extracted using a phenol-chloroform based method, as previously described^61^, and resulting DNA extracts were purified using a Zymo DNA Clean & Concentrator kit. Metagenome libraries were generated with a NEBNext Ultra II FS DNA library preparation kit, in accordance with manufacturer’s protocols. DNA fragment sizes were estimated using an Agilent Bioanalyzer DNA 7500 instrument with High Sensitivity kits depending on DNA concentrations and recommendations of protocols (Table S1). Sequencing of the 32 samples was performed at the Core DNA Sequencing Facility of the Fritz Lipmann Institute in Jena, Germany using an Illumina NextSeq 500 system (2 × 150bp). Resulting metagenomic library sizes ranged from 16.4 to 22.1 Gbp (mean = 19.6 Gbp; Table S1), and raw data was deposited into the ENA under project PRJEB36523.

### Metagenomic assembly and binning

Adapters were trimmed and raw sequences subjected to quality control processing using BBduk v38.51 ^62^. Assembly and binning were performed as previously described ^63^. Briefly, all libraries were independently assembled into scaffolds using metaSPAdes v3.12 ^64^, all of which were taxonomically classified per Bornemann et al. ^63^. For individual assemblies, open reading frames (ORFs) were identified using Prodigal v2.6.3 in meta mode ^65^. To generate coverage profiles, all quality-assessed and quality-controlled (QAQC) sequences from each of the 32 metagenomic libraries were mapped back to each of the 32 scaffold databases using Bowtie2 v2.3.4.3 in the sensitive mode ^66^.

Scaffolds were binned using differential coverages and tetranucleotide frequencies with Maxbin2 ^67^. Additionally, ESOM and abawaca ^68^ were used for both manual and automatic binning, based on tetranucleotide sequence signatures, using 3 kbp and 5 kbp or 5 kbp and 10 kbp as minimum scaffold sizes, respectively. DAS Tool ^69^ was used with default parameters to reconcile resulting bin sets. Complete sets of bins from each of the samples were dereplicated using dRep v2.4.0 ^70^. All scaffolds, bin assignments, ORF predictions, and taxonomic annotations were then imported into Anvi’o v6.0 ^71^. Each of the resulting 1,275 bins was manually curated in Anvi’o v6, considering both coverage and sequence compositions. In the end, 1,224 bins passed the 30% completeness [median = 61%, IQR = (49%,73%)] and 10% redundancy [median = 0%, IQR = (0%,1.4%)] quality thresholds.

### Characterizations of the Metagenome-assembled Genomes

ORFs originating from all of the resulting metagenome-assembled genomes (MAGs) were annotated using kofamscan ^72^ with the “detail” flag, and KO annotations were filtered using a custom script (https://git.io/JtHVw). This utility preserves hits with scores of at least 80% of the kofamscan defined threshold, as well as those exhibiting a score > 100 if there is no threshold. We elected to relax the default thresholds since all MAGs representing putatively chemolithoautotrophic microbes were verified manually, and we noticed that the best reciprocal blast hits with known reference sequences routinely scored below the kofamscan thresholds, *i*.*e*., we favored false positives over false negatives since we included a secondary verification step.

KEGGDecoder ^73^ was used to assess the metabolic potential of five of the primary chemolithoautotrophic pathways: the Calvin-Benson-Bassham cycle, the Wood-Ljundahl pathway, the reverse citric acid cycle, the 4-hydroxybutyrate 3-hydroxypropionate pathway, and the 3-hydroxypropionate bicycle. MAGs were examined in greater depth if a given pathway was > 50% complete. MAGs representing potential chemolithoautotrophs were re-annotated using the online BlastKoala server ^74^ with essential steps verified through blast ^75^ against the RefSeq database. A collection of HMM models was used to determine which form of Rubisco was detected, along with potential hydrogenases ^41^. Using blastp ^75^, dissimilatory bisulfite reductases (dsrAB) were compared to a database compiled by Pelikan *et al*. ^76^ to predict whether the pathway operated in an oxidative or reductive manner. Blast was used to compare gene hits for narGH/nxrAB (nitrate reductase / nitrite oxidoreductase) to a custom database based on sequences presented within Lücker *et al*.^77^.

All QAQC reads were remapped to a database consisting of only contigs of dereplicated MAGs. Normalized coverages for each of the MAGs was determined by scaling the resulting Anvi’o-determined coverages based on the number of RNA polymerase B (*rpoB*) genes identified in the QAQC-filtered reads. *RpoB* sequences were identified using ROCker with the precomputed model ^78^. Scaling factors were calculated by dividing the maximum number of *rpoB* identified in the 32 metagenomic libraries by the number of *rpoB* detected in each sample. Reported values represent averages of the triplicates/replicates, unless stated otherwise. The taxonomy of each MAG was evaluated using the GTDB_TK tool kit ^79^ in concert with the Genome Taxonomy Database (release 89) ^80,81^ and its associated utilities ^65,82–86^. Single copy marker genes were identified and aligned bwith GTDB_TK for all bacterial MAGs, and a phylogenetic tree of the concatenated alignment was constucted using FastTree2 v2.1.10 in accordance with the JTT+CAT evolutionary model. The resulting phylogenetic tree was then imported into iToL ^87^ for visualization, and all MAGs were subjected to growth rate index (GriD) analysis within each metagenomic library^88^.

Previously generated mRNA-enriched and post-processed metatranscriptomic libraries were procured from project PRJEB28783^37^. The groundwater source of these metatranscriptomes was collected in August and November 2015. QAQC filtered reads were mapped to MAGs using Bowtie2 v2.3.5 in sensitive mode^66^, and the total number of *rpoB* transcripts from each metatranscriptomic library were determined, as described above for metagenomes. The transcriptomic coverages for each ORF from each MAG were determined using Anvi’o v6 and normalized via scaling factor calculations based on the total number of *rpoB* reads from the original metatranscriptome library (*i*.*e*., the coverage of each ORF from each MAG was normalized to a community-wide estimate of the transcriptional activity of a house-keeping gene in each sample). Means were determined considering all of the metatranscriptomes generated from a given well, including different sampling timepoints. While well H32 was only sampled once, mean values from all other wells account for three to four metatranscriptome coverages each. Additionally, an average of the resulting normalized coverages for each MAG from each sample (sum of the MAG transcriptional coverage divided by the number of ORFs) was determined to estimate the relative transcriptional activity of the MAGs across the transect. Data was compiled and processed using R v.3.5.2 with Rstudio v1.1.463^89,90^ and the tidyverse package^91^, and color schemes were generated using the RColorBrewer utility^92^. All MAGs were deposited in project PRJEB36505’s data repository.

## Acknowledgments

We thank Falko Gutmann, Heiko Minkmar, Perla Abigail Figueroa□Gonzalez, and the Hainich CZE site manager Robert Lehmann for their assistance with sample preparation, collection, and filtration. We also thank Martin Nowak and John Southon for advice regarding ^14^C-bicarbonate supplementation and quantification of AMS results, respectively. Additionally, we thank Ivonne Görlich and Marco Groth from the Core Facility DNA sequencing of the Leibniz Institute on Aging - Fritz Lipmann Institute in Jena for their help with Illumina sequencing. study is part of the Collaborative Research Centre AquaDiva of the Friedrich Schiller University Jena, funded by the Deutsche Forschungsgemeinschaft (DFG, German Research Foundation) – SFB 1076 –Project Number 218627073. ST, XX and VFS acknowledge additional support from the European Research Council (Horizon 2020 Research and Innovation Programme, grant agreement 695101). MT gratefully acknowledges funding from the DFG under Germany’s Excellence Strategy - EXC 2051 - Project-ID 390713860. TLVB and AJP were supported by the Ministerium für Kultur und Wissenschaft des Landes Nordrhein-Westfalen (‘Nachwuchsgruppe Dr. Alexander Probst’). Climate chambers to conduct experiments under controlled temperature conditions were financially supported by the Thüringer Ministerium für Wirtschaft, Wissenschaft und Digitale Gesellschaft (TMWWDG; project B 715-09075). The scientific results have in part been computed at the High-Performance Computing (HPC) Cluster EVE, a joint effort of both the Helmholtz Centre for Environmental Research - UFZ (http://www.ufz.de/) and the German Centre for Integrative Biodiversity Research (iDiv) Halle-Jena-Leipzig (http://www.idiv-biodiversity.de/).

## Data Availability

The metagenomic raw data, individual sample assemblies, and the metagenome-assembled-genomes for this study were deposited into the ENA under project PRJEB36523. All raw and summarized AMS data is available in the Supplemental Data File.

## Author contributions

KK, ST, WAO and MT designed this study. WAO, KK, AJP, TLVB, MM, MH, and MT planned, designed, and conducted the metagenomic sampling approach. WAO, TLVB, and AJP performed the metagenomic analysis. WAO, ST, XX, and VFS performed the carbon fixation experiments and interpreted the results. MK, MH, BT, and LB conducted the anammox and nitrification rate measurements. WAO, KK and ST wrote the manuscript with the help of all authors.

## Figures

(Vector graphics versions of all figures are attached as a separate document)

## Notes

### Competing Interest Statement

The authors have declared no competing interest.

