## Supplemental Information for "Rates of primary production in groundwater rival those in oligotrophic marine systems"

### Quantification of carbon fixation rate under *in situ* conditions

The experimental setup included two types of controls, kill controls that were amended with the same level of $^{14}C$-bicarbonate along with 37 mmol $ZnCl_2$ to inhibit microbial metabolism and no-label addition controls. The no-label controls always yield lower 14C/12C ratios than the kill controls indicating some label carry over or incomplete inhibition. This translated to 3.7% to 17.7% higher rate measurements using the no-label controls over the kill controls (Table S2). In one instance, well H51, the kill controls had higher 14C/12C values than the corresponding labeled bottles. This sample was removed from the main analysis. However, in Figure S4 we estimated an expected $CO_2$ fixation rate using the rate based on the no-label addition controls that was corrected using the maximum difference in rate measurements between the kill controls and no-label added controls for the other 5 wells (17.0%).
The error estimations presented in the main text and Table S2 utilize POC measurements from all 9 bottles (3x labeled, 3x kill controls, 3x no-label controls).

### Conservative estimations of net primary productivity within the groundwater

The use of $^{14}C$-bicarbonate additions to estimate carbon fixation rates in marine systems is thought to approximate net primary production rather than gross primary production, and is further thought to underestimate these net production rates[1,2]. The quartz fiber filters (QFFs) used here, with a nominal pore size of 0.3 µm, were chosen to withstand the 900 □ baking temperature. However, they likely missed a portion of the ultra-small microorganisms that are abundant within groundwater[3], and this aquifer in particular[4], contributing to our expectation of conservative estimations. It is also likely the organic carbon retained within the QFFs filters in part contained adsorbed DOC, as has been observed in the marine habitats[2]. If this adsorbed DOC was primarily old carbon, as indicated by previously measured $^{14}C$-DOC signatures[5], our rate calculations would further underestimate true values.

Furthermore, our primary production rates only consider the planktonic portion of the community, and previous studies have suggested a portion of the autotrophic community is attached within this aquifer system[6].

### Carbon fixation rate variation within the transect

Within our aquifer transect, a one-way analysis of variance (ANOVA) revealed significant differences in carbon fixation rate between the wells ($F(14) = 6.7$, $p < 0.007$; Figure 1 A). There were significant differences between the two wells with the highest rates (H52, H43) and the two wells with the lowest rates (H14, H41) ($p<0.05$). H32 was not significantly different from any of the other wells, with means falling between the two extremes. The transect locations used in this study were chosen to best represent all the identified biogeochemical regimes and each represents a distinct microbial community[7]. Indeed, previous work by Yan et al., has identified a complex interplay of hydrochemical parameters that could explain microbial variation, including ammonium, nitrate, dissolved oxygen, reductive potential, and Fe(II). The transect wide carbon fixation rates identified here were significantly related to redox potential, and were higher in wells with the lowest redox potentials ($F(1)=22.2$, $p=0.0004$), but with the relatively small dataset were not significantly correlated to dissolved oxygen concentrations (Figure 1 B, C).

### Distributions and relative abundances of putatively chemolithoautotrophic MAGs

The 102 putatively chemolithoautotrophic MAGs were significantly more enriched in the 0.2 μm filter fraction (mean normalized coverage = 13.6, median = 1.4) compared to the 0.1 μm fraction (mean = 3.1, median = 0.25) (Wilcoxon paired test, V=115900, $p < 2.2e-16$). Based on the percentage of QAQC reads that mapped back to each of the MAGs, they collectively represented 0.8% to 11.5% of the total sequenced microbial community in the 0.2 μm fraction, compared to 5.5% - 36.1% of the reads mapped to the other 1122 MAGs within the same fraction (Figure S3). Using the more appropriate, but less intuitive, normalized coverage scale that is not influenced by genome size, the sum of all chemolithoautotrophic MAGs ranged from 225-2327 across the transect within the 0.2 μm fraction compared to 1157-10691 for the others within the same filter fraction (Figure S3). These chemolithoautotrophic MAGs had significantly higher growth rate indexes (GRiD scores, median = 1.79 IQR=[1.47, 2.21]) compared to the other MAGs (median = 1.52, IQR=[1.27, 1.86], Wilcoxon test, V=814730, $p<2.2e-15$).

### Patterns in MAG electron donor sources

Inferring the likely electron donor pathways based on the metabolic potentials of the representative MAGs suggests that reduced nitrogen is fueling nitrification within H41 and plays a lesser role in H51 and H32 (Figure 4B). Based on summed coverages, reduced sulfur appears to be driving carbon fixation with H52, but is also important in H32, and H51. The potential for anaerobic ammonium oxidation coupled to the Wood Ljungdahl pathway within Planctomycetota MAGs was also highest in H52, but these MAGs were not as relatively abundant as the reduced sulfur and CBB utilizing Proteobacteria. However, there was a similar recruitment of transcripts within these groups (Figure 3). This matched results previously shown by Wegner and colleagues[8] that genes involved in anaerobic ammonium oxidation had a high ratio of transcripts to DNA and indicating that anammox may play a larger role in carbon fixation within the anoxic groundwater of H52.

### Mismatch in $CO_2$ fixation and chemolithoautotrophic potentials

Wells H43 and H52 had the highest rates of carbon fixation within the aquifer transect. Both were characterized by lower redox potentials (~250 mv) and mostly anoxic conditions (H52 was always anoxic, while H43 periodically exhibited trace oxygen levels). However, the wells are consistently grouped into distinct geochemical clusters[9] and have a distinct microbial community[7]. The high rate within H52 matched previous hypotheses about the importance of chemolithoautotrophy within this well[8,10]. In contrast, H43 did not match our expectations of a low rate of carbon fixation based on previous estimates for the importance of chemolithoautotrophy[10]. Additionally, in our study, H43 had among the lowest relative abundances of potential chemolithoautotrophic MAGs - most of which did not have genetic evidence for the acquisition of electron donors (Figure 1 C).

At this time, we do not have a compelling explanation for these contrasting patterns. The DNA samples for the metagenomes and the water collected for the rate measurements did come from different sampling campaigns. However, the microbial communities appear stable throughout the year and over longer interannual cycles[7] which does not support the hypothesis that H43 had a lower chemolithoautotrophic potential only during the DNA collection time point for this study. Total iron content is highest in well H43 and previously identified as an explanatory variable differentiating anoxic groundwater wells[11]. We postulated that iron-oxidizers may be more important in this region and subsequently screened all all MAGs for known iron-oxidizing functions[12]. Results were inconclusive, likely due to paucity of information about mechanisms of iron-oxidation[13,14]. It also may be that more oxygen was introduced into the samples collected from H43, providing extra electron acceptor and generating more of a potential carbon fixation rate than what was observed *in situ*. We do not favor this explanation since the replicates for H43 are within the range of the H52 samples.

We also hypothesized that H41 would exhibit high rates of carbon fixation, due to a preponderance of nitrifying chemolithoautotrophs, found in this study and by Wegner et al., 2019. Normalizing the rate based on expected bacterial populations sizes, does help explain the difference in absolute rates (Table S5). However, based on the standing concentration of particulate organic carbon within the different regions, H41 primary production contributes among the lowest amount. The ammonium concentrations within H41 are nearly 10 µM, much lower than the 35 and 27 µM concentrations seen in H52, and H43 respectively[10], but unlikely to be limiting for the carbon fixation rate measured (>200x higher than necessary). The $^{14}C$ signatures of dissolved organic carbon within H41 indicated that younger surface derived carbon was more prevalent in this well, H51, and H32 than in H43 and H52 [5]. H41 also exhibits the largest variation in $^{14}C$-DOC signatures, with an influx of younger dissolved organic carbon in April - July, when our $CO_2$-fixation samples were collected[5]. Therefore, carbon fixation may play a less important role during these months.

### Metabolic potentials and activities of abundant primary producers within anoxic groundwater

Next we focused on the most abundant MAGs within the groundwater with the highest rates of carbon fixation to understand specific metabolic features of the most important chemolithoautotrophic populations. In general, these MAGs tended to be detected under a wide range of biogeochemical conditions (Figure 3). MAG H51-bin250-1, belonging to the class Nitrospiria, was the most relatively abundant MAG across all wells, particularly under anoxic and hypoxic conditions (Figure 3). Classified within the order “9FT-COMBO-42-15”, this MAG was 91% complete with an estimated 1.4% redundancy. There was a complete carbonyl branch of the Wood-Ljungdahl pathway, and a partial methyl pathway containing folD and metF, but missing formate-tetrahydrofolate ligase (fhs) and formate dehydrogenase (fdhAB). By incorporating previous RNA-Seq data[8], we were able to assign mRNA transcripts to all genes from both branches (Figure 4 A). Reducing energy could be provided by forms of sulfur, using either the oxidative dissimilatory sulfate reduction pathway (the reverse Dsr pathway), or thiosulfate through the presence of a partial SOX pathway (soxBY)[15]. There were three potential electron accepting processes, oxygen through a cytochrome c oxidase, and nitrate, either through dissimilatory reduction to ammonia (DNRA: narGHI, nrfA, and nirB) or via denitrification (nirS, norC, nosZ, but missing norB). Only transcripts to genes involved in nitrate reduction were recruited, even in wells with oxygen present (Figure 4 A). The presence of multiple terminal electron accepting processes other than $CO_2$ suggest a mixotrophic lifestyle, similar to other Wood Ljungdahl utilizing acetogens[16,17]. There is precedence for autotrophic WL-utilizing bacteria from Nitrospirota, although the only example, *Ca.* Magnetobacterium, is found within class Thermodesulfovibrionia[18]. *Ca.* Magnetobacterium was also described with a similarly flexible metabolism with the capacity for sulfur oxidation and denitrification, suggesting these traits may be more widespread within the phylum.

Proteobacteria within the GTDB-named family Sulfurifustaceae (Figure 3, node 10) were also relatively abundant within well H52, and to a lesser extent H32. The most abundant representative chemolithoautotrophic MAGs were H32-bin014 and H32-bin069, neither of which were classified below the family level. These two MAGs had almost identical distribution patterns and transcript recruitment numbers. Both exhibited lower completion values at 38% and 43% respectively, and each was estimated to be 1.4% redundant. They shared 94.5% amino acid identities (AAI), and appeared to fill a similar ecological niche. Both included copies of form II rubisco, typical for Proteobacteria from low $O_2$ and high $CO_2$ environments[19]. Important pathways for nitrogen cycling were more complete within H32-bin014, while H32-bin069 contained more complete pathways for sulfur cycling processes (Figure S5). They showed a large flexibility in electron acceptors including both cytochrome c oxidase (complex IV) and the high $O_2$ affinity *cbb3* type cytochrome c oxidase common to Proteobacteria[20], along with genes involved in DNRA. Electrons are expected to be provided by reduced sulfur sources, via the oxidative Dsr pathway and the Sox pathway, which are both highly expressed (Figure 4, Figure S5). In support of flexible energy conserving mechanisms, both recruited transcripts from wells that spanned the full oxygen gradient within the transect. While the normalized coverage of H32-bin014 within H52 is lower than the Nitrospiria MAG H51-bin250-1 (130x vs 201x), the transcription of genes involved in carbon fixation and energy conservation were >10x higher (Figure 3).

The closest reference genomes were *Sulfuricaulis limicola* (68% AAI) and a Ca. Muproteobacteria labeled RIFCSPHIGHO2_12_FULL_60_33 (67% AAI). Under the GTDB classification nomenclature, Muproteobacteria fall within the Sulfurifustaceae family, and this group has been linked in both aquatic and terrestrial environments for the potential to oxidize sulfur using both oxidative Dsr and Sox pathways[21,22]. *S. limicola* was isolated from freshwater sediment and was experimentally shown to use thiosulfate, tetrathionate, and elemental sulfur as electron donors to fuel CBB-based carbon fixation[23]. Complete genome sequences found the presence of both oxidative Dsr and Sox pathways[24].

While much less relatively abundant, Planctomycetota MAGs within the family Brocadeiaceae exhibited a similar mean transcriptional activity score as the Sulfurifustaceae MAGs (Figure 3, node 20). H41-bin243-1 was one of the more complete representatives (85%), with no evidence of redundancy, and was among the most transcriptionally active MAGs within this phylum (Figure 4 C). The products of the most transcribed genes were similar to proteins involved in the anaerobic oxidation of ammonium (anammox), particularly hydrazine dehydrogenase (Hdh - K20935), and one of the six copies of hydroxylamine oxidoreductase

(Hao - K10535). These recruited approximately 10-fold more transcripts than the genes for carbon fixation (Figure 4 C). Within anammox granules grown in a bioreactor, these two specific functions were also the most highly expressed and were 10-20 fold higher than the median transcriptional rate[25], and genomes from this group are known to contain a large number of orthologs to Hao[26].

The most transcriptionally active MAGs (H51-bin43-1, H51-bin207-2, H41-bin278-1, H52-bin010-1, H52-bin125-1) fell within a tight phylogenetic cluster, sister to a clade formed by *Ca.* Jettenia caeni, *Ca.* Brocadia fulgida, and MAG H32-bin113-1 (Figure S6). The relatively more abundant but less transcript-recruiting MAG, H41-bin245-1, was closely related to *Ca.* Scalindua rubra-A (Figure 3 node 21, Figure S5, AAI = 74%). The transcriptional activity within this diverse clade of putative anammox bacteria suggested that they play a disproportionately large role in chemolithoautotrophy relative to their DNA-based abundances. Surprisingly, all putative anammox MAGs were detected and transcriptionally active in the fully oxic groundwater from H41 and H51 (Figure 3, Figure 4 B). Anammox is typically reversibly inhibited in the presence of oxygen[27], although under low oxygen environments these microorganisms will still express the necessary functions[28,29]. However, anammox bacteria can be protected from higher oxygen concentrations by microbial biofilms, and under bioreactor conditions the highest rates of nitrogen removal have coincided with 3 mg $L^{-1}$ DO[30]. The metagenomic samples libraries were collected from ~100 l of groundwater, while the metatranscriptomic libraries were generated from ~2000 l, which perhaps was more disruptive to intact biofilms and explains greater enrichment in the metatranscriptomes.

Widely distributed across the transect were 10 putative WL using chemolithoautotrophs, all within the class Thermodesulfovibrio (Figure 3). None were affiliated to the same family as *Ca.* Magnetobacterium, and while the WL pathways were mostly complete, all the MAGs contained *dsrAB* genes affiliated with the canonical reductive pathway, rather than the oxidative pathway, as seen in H51-bin250-1. Similar to the deep-branching Thermodesulfovibrio MAG described by Arshad et al. [31] many contained genes associated with DNRA or denitrification. The low relative abundances and lower levels of transcripts mapping to this group suggests that even if they were fixing carbon through the WL pathway they may not play a large role in organic carbon production within the aquifer transect. The lack of pathway to access inorganic electron donors suggests they may grow heterotrophically, utilizing the WL-pathway to completely oxidize acetate as seen in some sulfur reducing bacteria[32].

Dominant MAGs within oxic groundwater

### Metabolic potentials and activities of abundant primary producers within oxic groundwater

The most relatively abundant chemolithoautotrophs within oxygen containing groundwater also belong to class Nitrospiria (Figure 3, Figure S3). Encoding the rTCA cycle, they fell within two distinct functional clades, complete ammonium oxidizing bacteria (comammox) and nitrite oxidizing bacteria. The comammox groups were more relatively abundant, and are represented by two phylogenetically distinct clades including the MAG H41-bin216-1 (genus *Nitrospira*), and MAGs H51-bin251-1/H41-bin006-1 (genus Palsa-1315). Based on KEGG annotations, these two clades of comammox MAGs shared a similar metabolic profile, with some of the only differences being that the Palsa-1315 MAGs had superoxide dismutase and catalase genes (SOD/CAT) which were lacking in the *Nitrospira sp*. Their metabolic potential was largely congruent with previously published genomes of comammox bacteria[33]. Here, the best representatives from both clades had the same completeness of the rTCA cycle, including ATP citrate lyase (aclAB), pyruvate:ferredoxin oxidoreductase (POR), and fumarate reductase (FRD), and both were missing all four ORFs associated with 2-oxoglutarate:ferredoxin oxidoreductase (korABCD). They also contained genes for ammonium monooxygenase (amoABC), hydroxylamine oxidoreductase (hao), and nitrite oxidoreductase (nxrAB), along with NO-forming nitrite reductase (nirK) and NADH-dependent nitrite reductase (nirD). They contained mostly complete flagella and chemotaxis associated proteins. The normalized transcriptomic patterns were largely congruent with the DNA-based normalized coverages (Figure 3). The MAGs recruited among the highest number of transcripts from the H41 well containing high oxygen concentrations.

The two distinct clades of putatively nitrite oxidizing Nitrospiraceae MAGs, were less relatively abundant than the comammox MAGs and were best represented by H41-bin184-1 (genus *Nitrospira_D*) and H41-bin274-1 (*Nitrospira_A*) (Figure 3). Their distributions (both DNA-based normalized coverages and normalized transcripts) overlapped with the related commamox Nitrospiraceae MAGs and were also only missing the *korABCD* ORFs from the rTCA cycle. A third putative nitrite oxidizing MAG (H14-bin041-1) was only classified to the order level (Nitrospirales), with a very different distribution that skewed towards wells with low, but detectable oxygen levels (H32, H51, H14). This MAG was only 65% complete, with no detected redundancy, but the nxrA gene (nitrite oxidoreductase) was most similar (76%) to the two reference nxrA gene copies found in *Nitrospira defluvii*[34].

Proteobacterial chemolithoautotrophs utilizing the Calvin Benson Bassham (CBB) cycle were also well represented under oxic conditions (Figure 3). The most relatively abundant and active proteobacterial MAGs in the oxic wells were predicted to use reduced nitrogen to conserve energy. There were two clusters of very closely related MAGs that fell within the Nitrosomonadaceae family (Burkholderiales). MAGs H41-bin218-1, H51-bin262-1, and H51-bin202-1 were within the *Nitrosomonas* genus and were >97% average amino acid identity to each other. The three MAGs each recruited among the the most transcripts of any of the other MAGs we recovered (Figure 3). The most complete MAG, H41-bin218-1 was estimated to be 94% complete and 0% redundant (Table S3) and had a mostly complete CBB cycle with form I rubisco ( 2 copies of both large and small subunits), glyceraldehyde 3-phosphate dehydrogenase, phosphoribulosekinase, transketolase, ribulose-phosphate 3-epimerase, and ribulose-5-phosphate isomerase, similar to the *N. europaea* genome[35]. The ammonium oxidation pathway was less complete, containing a complete amoC gene, a partial amoA gene, and missing amoB. However, the very closely related H51-bin202-1 had all three genes, and the cluster of genomes from which H41-bin218-1 was selected from also contained all three genes. These representative MAGs were also missing hydroxylamine oxidoreductase (Hao), but again, the larger genome clusters for each did contain high confidence hits for Hao. Also similar to *N. europaea*, there were copies of nitrite reductase (nirK), but they were missing nitric oxide reductase (norBC). These MAGs were predicted to be motile with complete flagellar synthesis genes and chemotaxis genes.

The second cluster of H51-bin263-1, H41-bin193-1, and H41-bin007-1 showed approximately the same summed relative normalized abundance values but were more abundant in H51 than in H41. Transcripts mapping to these MAGs were from both wells, with higher normalized numbers mapping from H41 than H51. The best representative was H51-bin263-1 with 93% completeness and 0% redundancy estimates. Similar to the *Nitrosomonas* cluster, H51-bin263-1 had a mostly complete CBB cycle with the same genes present. Unfortunately, this particular representative only had nirK and copies for amoABC and Hao were only found within the larger genome cluster (H41-bin007-1 did have a amoC copy).

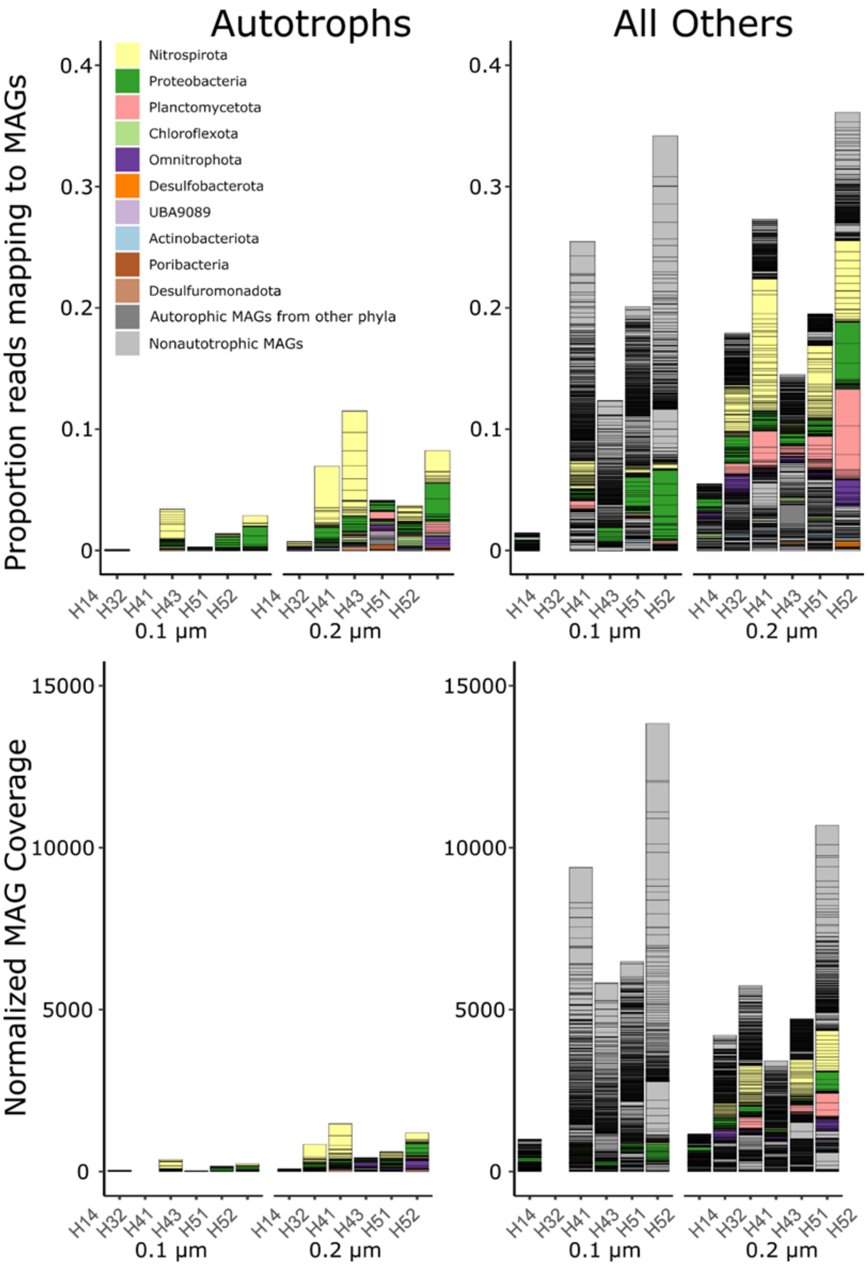


Figure S1. (Top) Proportion of metagenomic short reads that mapped to the putatively chemolithoautotrophic MAGs. Colors indicate the phylum and each bar indicates a specific MAG. (Bottom) The normalized coverages assigned to the MAGs, which unlike the proportional representation is independent of MAG size. For each plot, 0.1 µm and 0.2 µm represent the filter fraction from which the DNA was extracted from.

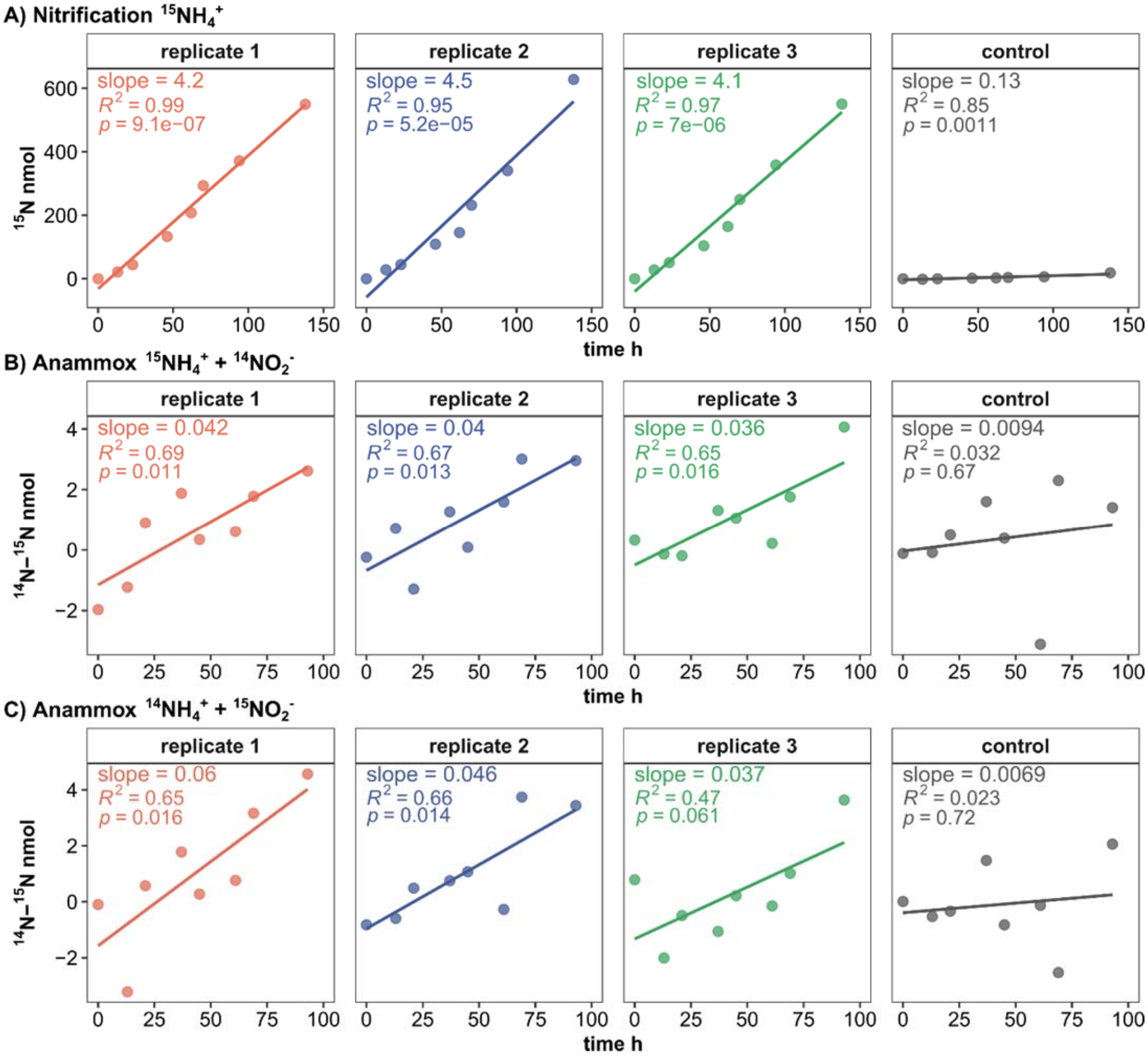


Figure S2. Assessment of nitrogen transformation process rates in groundwater incubations by linear regression of measured increase in $^{15}N$ over time for (A) nitrification with $^{15}NH_4^+$ label, (B) anammox with $^{15}NH_4^+$ label and (C) anammox with $^{15}NO_2^-$ label. Shown are the slopes of the regression line for each replicate and control sample from which nitrification and anammox rates were calculated. $R^2$ values describe the accuracy of the linearity and *p*-values the significance of results for $p < 0.05$.

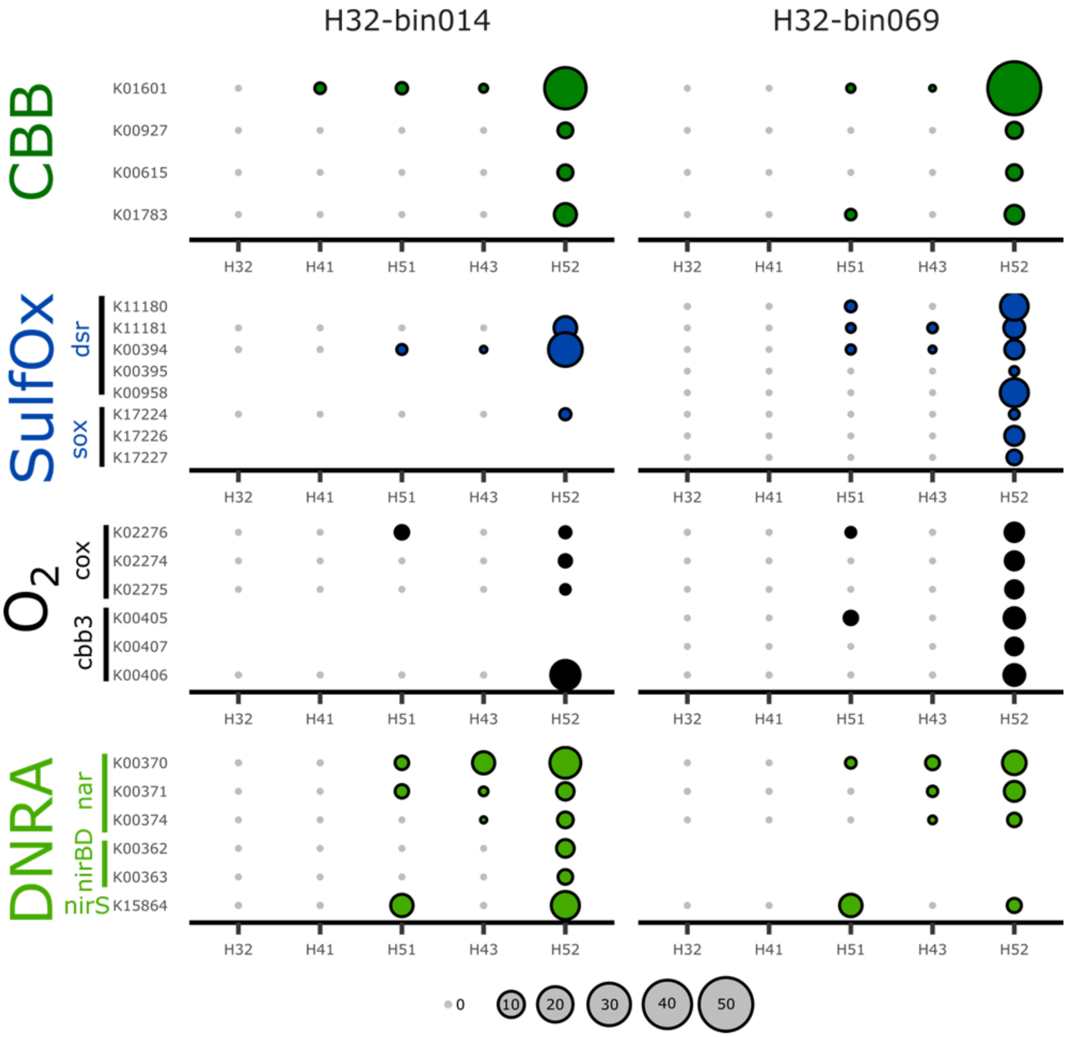


Figure S3. Transcriptional activity of both dominant MAGs within the family Sulfurifustaceae, exhibiting nearly identical distributions throughout the aquifer transect and a >94% AAI. H32-bin014 is also shown in Figure 5 B. Bubble size is relative to the normalized ORF transcriptional coverage.

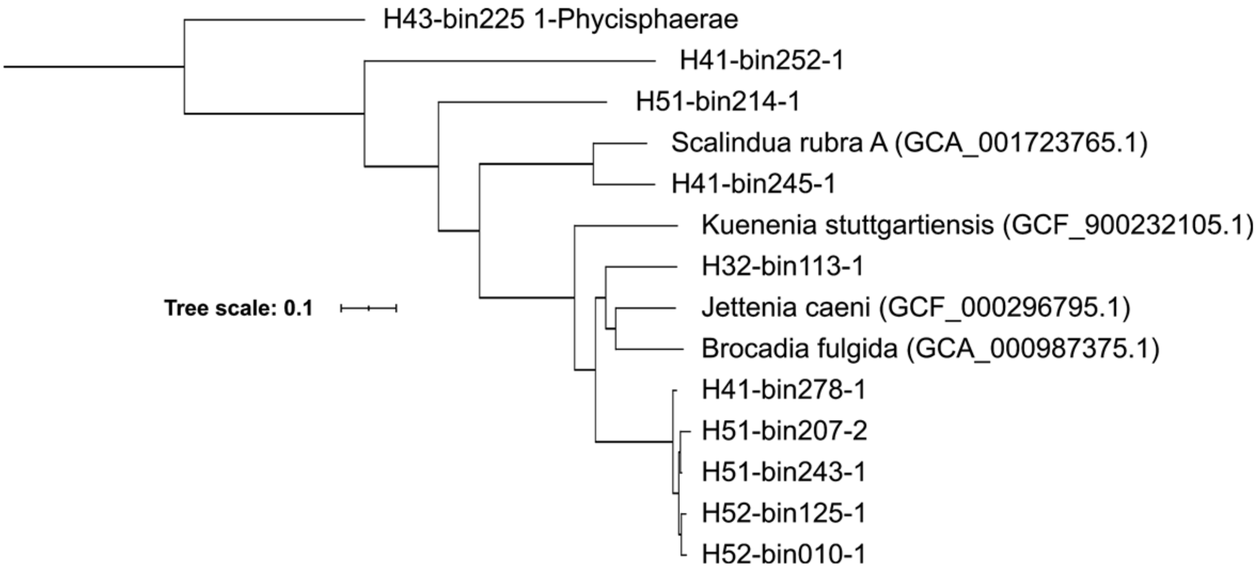


Figure S4. Approximately maximum-likelihood phylogenetic tree of Wood Ljungdahl containing anammox MAGs and reference genomes. The multiple protein sequence alignment from GTDB_TK (5040 positions) was used with the JTT+CAT model, manually rooted with a MAG classified to the sister class Phycisphaerae within the phylum Planctomycetota.
